# Interoperable genome annotation with GBOL, an extendable infrastructure for functional data mining

**DOI:** 10.1101/184747

**Authors:** Jesse C.J. van Dam, Jasper J. Koehorst, Jon Olav Vik, Peter J. Schaap, Maria Suarez-Diez

## Abstract

**Background:** A standard structured format is used by the public sequence databases to present genome annotations. A prerequisite for a direct functional comparison is consistent annotation of the genetic elements with evidence statements. However, the current format provides limited support for data mining, hampering comparative analyses at large scale.

**Results:** The provenance of a genome annotation describes the contextual details and derivation history of the process that resulted in the annotation. To enable interoperability of genome annotations, we have developed the Genome Biology Ontology Language (GBOL) and associated infrastructure (GBOL stack). GBOL is provenance aware and thus provides a consistent representation of functional genome annotations linked to the provenance. GBOL is modular in design, extendible and linked to existing ontologies. The GBOL stack of supporting tools enforces consistency within and between the GBOL definitions in the ontology (OWL) and the Shape Expressions (ShEx) language describing the graph structure. Modules have been developed to serialize the linked data (RDF) and to generate a plain text format files.

**Conclusion:** The main rationale for applying formalized information models is to improve the exchange of information. GBOL uses and extends current ontologies to provide a formal representation of genomic entities, along with their properties and relations. The deliberate integration of data provenance in the ontology enables review of automatically obtained genome annotations at a large scale. The GBOL stack facilitates consistent usage of the ontology.

## Background

Advances in sequencing technologies have turned genomics into a data-rich scientific discipline in which the total assembled and subsequently annotated sequence data doubles every 30 months (1). To support the growth in data throughput, automated annotation algorithms have become an indispensable supplement to manual annotation (2, 3) and currently, automatic annotations in the UniProt database outnumber manual annotations 100-fold (4).

Functional genome comparison has been used to identify diagnostic markers, to develop effective treatments, and to understand genotype-phenotype associations (5–7). The volume and heterogeneity of genome annotation data has created a unique type of big data challenge, namely how to transform computational predicted annotations into actionable knowledge. Tapping into these available resources is only efficiently done by computational means and requires a consistent interlinking of data so that data becomes Findable, Accessible, Interoperable and Reusable (FAIR) (8).

The format for sharing of public genome sequence annotation data has been developed and is maintained by the International Nucleotide Sequence Database Collaboration (INSDC) a long-standing foundational initiative that operates between the DDBJ, EMBL-EBI and NCBI public repositories. However, tradeoffs between simplicity, human readability and representational power left little support for interoperability, i.e. the ability of computer systems to directly make use of information. The */inference* qualifier (9) provides a structured description of evidence that supports feature identification or assignment. Thus, within the standard formats, data provenance of computational annotations could be stored under this optional inference tag but this tag is not designed to be used for contextual, element-wise provenance.

Currently, most annotations rely on computational predictions of structure or function, and the choice of thresholds for confidence scores becomes a key consideration. Tracking the provenance of genome annotations becomes essential for scientific reproducibility and to enable critical reexamination of analyses. However, such meta-analysis is currently very time-consuming. Efficient meta-analysis would require a framework able to accommodate the various types of annotations (e.g. gene prediction, homology, protein domains) directly linked to the supporting statistical evidence. Presently, no machine-readable infrastructure exists to directly query genome annotations linked to the historical and contextual provenance. The World Wide Web consortium provides the Semantic Web and the Resource Description Framework (RDF) data model, supporting these requirements. For RDF, ontologies are essential as they provide consistency in the meaning of data elements and in the relationship between them (10).

In this respect, ontologies already exist for various aspects of biology (11). The Sequence Ontology (SO) (12) was presented over 12 years ago and was designed as a complete terminology of unambiguous terms related to genetics. However, it was never intended to function as a file format or database schema, and provides no support for linked sets of data attributes. Furthermore, it has limited support for storing based-on provenance except for some experimental codes. FALDO’s (13) only purpose is to unambiguously store genetic locations on a sequence. The Synthetic Biology Open Language (SBOL) (14) was successfully designed to describe complete synthetic constructs and the interactions between each of the elements. None of these standards were designed to consistently store feature predictions with evidence provenance and therefore none of these tools provides a complete representation of the genomic information linked to the provenance it is based on.

To meet the requirements and to ensure interoperability of computational predictions, we developed an extendable provenance-centered infrastructure for interoperable genome annotations. The here presented infrastructure consists of two main elements; Firstly, the Genome Biology Ontology Language (GBOL), which directly integrates evidence provenance for the whole dataset and for each included element (dataset- and element-wise provenance). Secondly, the "GBOL stack" of enforcing tools facilitates the consistent usage of ontologies. GBOL is modular in design, extendible and linked to existing ontologies. Empusa has been developed as part of the GBOL stack to ensure consistency within and between ontology (OWL), the API and the Shape Expressions (ShEx) describing the graph structure. This enables the use of SPARQL queries to include contextual details in large scale functional analyses. Modules have been developed to serialize the linked data (RDF) and to generate a plain text format files.

## Results

### Ontology structure

GBOL is a genome annotation ontology developed for the application of semantic web technologies in genome annotation and mining. As such GBOL provides the means to consistently describe computationally inferred genome annotations of biological objects typically found in a genome sequence annotation data file in the public repositories. Additionally, it can describe the linked data provenance of the extraction process of genetic information from genome sequences.

An overview of the structure of GBOL is shown in Figure 1. The ontology contains 251 classes that can be categorized into 6 broad domains (Table 1). In GBOL, sequences have features, which in turn have genomic locations on the sequence. The authority of this relationship is derived from the data provenance that captures both the statistical basis of each individual annotation (element-wise provenance) as well as the programs and parameters used for the complete set of sequences under study (dataset-wise provenance). All annotations for a given sequence can be packed into a single entity called a document.

**Fig. 1.**
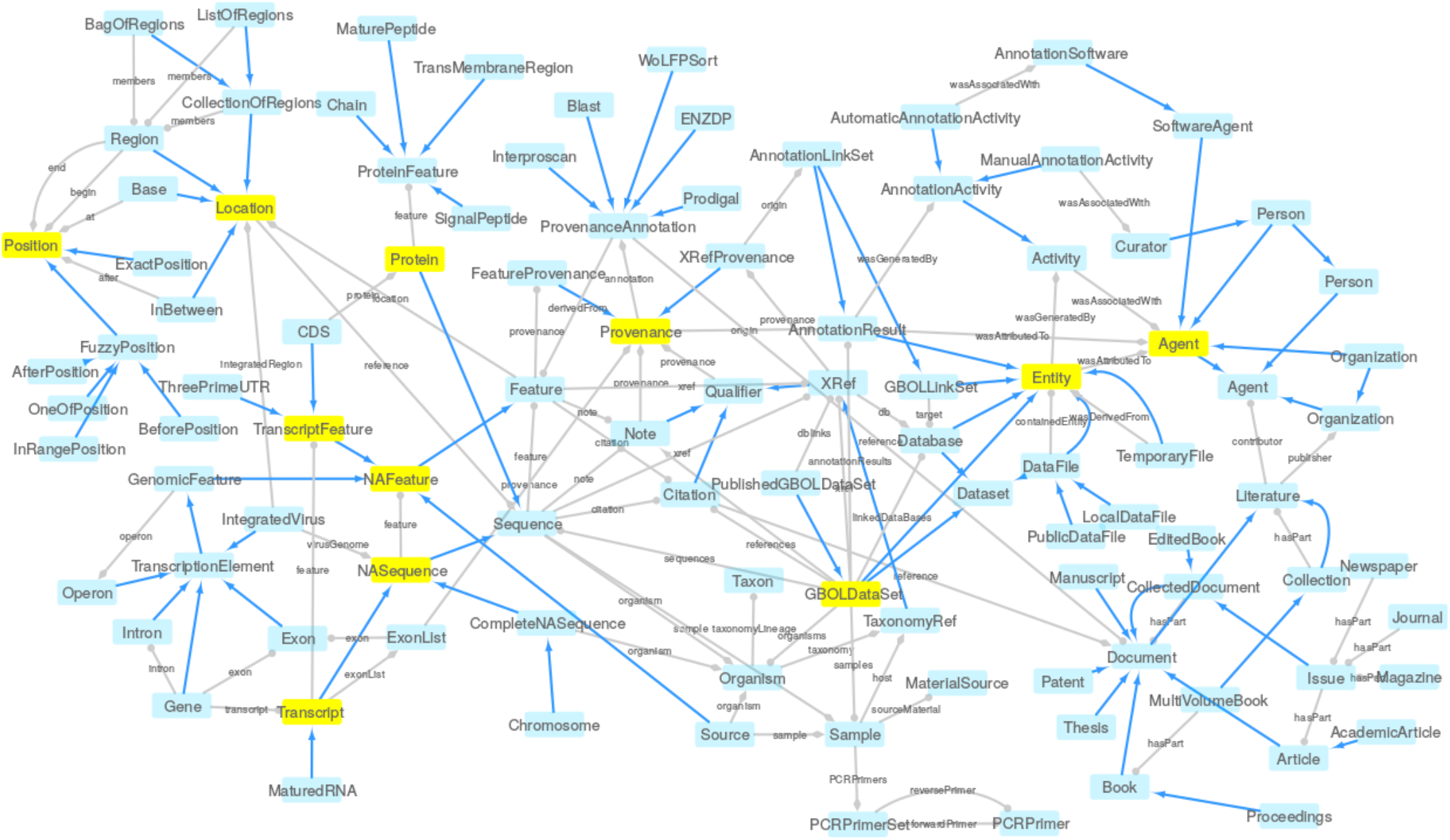
The GBOL ontology structure. Network based view generated using RDF2Graph(15) the GBOL core ontology. Nodes represent types. Blue edges represent *subClassOf* relationships whereas grey edges represent unique type links. A unique type link is defined as a unique tuple: type of subject, predicate, (data)type of object. Arrow heads indicate the forward multiplicity of the unique type links: 0¨1 and 1¨1 multiplicities are indicated by diamonds; 0¨N and 1¨N multiplicities are indicated by circles. Neighbourhood of nodes marked in yellow is further expanded in Figures 4–8

**Table 1.** Overview of domains, classes and properties described by the the GBOL ontology. Note that some properties might be in multiple sub domains.

| Sub domain | Classes | Properties | Value sets |
| --- | --- | --- | --- |
| Genomic locations | 16 | 17 | 1 |
| Genes |  |  |  |
| transcripts and features | 114 | 133 | 17 |
| Document structure | 27 | 107 | 7 |
| Dataset-wise provenance | 22 | 54 | 0 |
| Element-wise provenance | 5 | 9 | 0 |
| BIBO | 59 | 90 | 2 |

### Design principles

GBOL was developed focussing on its function as as file format and as database schema and has the following design principles: modularity, human readability, and annotation. These principles ensure that the ontology can be easily extended (16).

#### Modularity

The number of classes in the main class tree is kept as small as possible and elements within the data are described with attributes when possible. Furthermore, classes are included in the main class tree only when there are unique properties in a class or in one of the sibling classes. This approach ensures that sub-ontologies can be managed as separate entities within the main ontology and that we can use existing ontologies. As an example the class *RegulationSite* has an attribute *regulatoryClass,* which denotes the type of regulation with a separate set of classes of which all are instances of the regulatory class.

To further simplify the ontology, every attribute is defined as a direct property within the class that links to either a string, an integer, another object or a class in an enumeration set. For each class in which the attribute is used, an ‘all values from’ axiom is used, with an optional minimal and/or maximal cardinality constraint. The ’all values from’ axiom enforces all referenced objects to be of the expected type, which is not the case with the ‘some values of’ axiom and therefore we excluded the use of the ’some values of’ axiom. This approach is fundamentally different from the principle used in the SO, in which attributes are defined using the ’has quality’ property in combination with the ‘some values of’ axiom that references to a class.

#### Human Readability

All names within the ontology adhere to a set of basic principles to increase (human) readability of the ontology. All class names represent the underlying biological concept as closely as possible avoiding the use of unreadable numbers. All classes start with uppercase whereas properties start with lowercase. All words are spelled out, and white spaces are left out of the names, instead the next word starts with uppercase. In this way, the class ‘exact position’ becomes ‘ExactPosition’ and the property ‘regulatory class’ becomes ‘regulatoryClass’. Furthermore, where possible, the names are shortened with abbreviations, as long as they remain understandable for a human reader (e.g. XRef instead of CrossReference).

#### Annotation

All classes and terms within the ontology are annotated with a short definition; an optional comment with additional usage information; an optional editorial comment relating to the development of the ontology itself; an optional ddbj label indicating the presence in the GenBank standard; and an optional SKOS (17) exact match to relate classes to terms in existing ontologies.

### The GBOL infrastructure

An infrastructure enabling interoperable genome annotations integrated with provenance requires the following characteristics: i) An OWL (18) encoded definition of an ontology. ii) An infrastructure to enhance and simplify its usage, consisting of an interface (API) that allows to use Java and R. iii) A file format that can be obtained from serializing the linked data (RDF) using a lightweight Linked Data format (JSON-LD) (19) which is subsequently serialized as YAML (20). This format mimics the layout of the current format for sharing of public genome sequence annotation data, but has integrated support to add additional information. iv) A ShEx definition for data conformance validation to enhance data consistency (21). And v) a tool to convert existing GenBank and EMBL format files into the GBOL format.

GBOL data can be stored in any of the linked data formats (RDF), such as Turtle. The generated API can be used to access the genomic information encoded within the GBOL format, which includes a data consistency validation. The API directly reads from and directly modifies the RDF data structure upon usage of any of the data model functions. This enables the usage of SPARQL within the client code, which can run a SPARQL query and directly use the resulting objects nodes in the API. Moreover, the RDF data can be structured into a tree with the JSON-LD framing API into JSON-LD, which, in turn, can be further serialized as YAML resulting in a human readable format for sharing of public genome sequence annotation data. By addition of standard annotation tools, the GBOL stack can be at the core of a provenance-centered genome annotation framework (Figure 2).

**Fig. 2.**
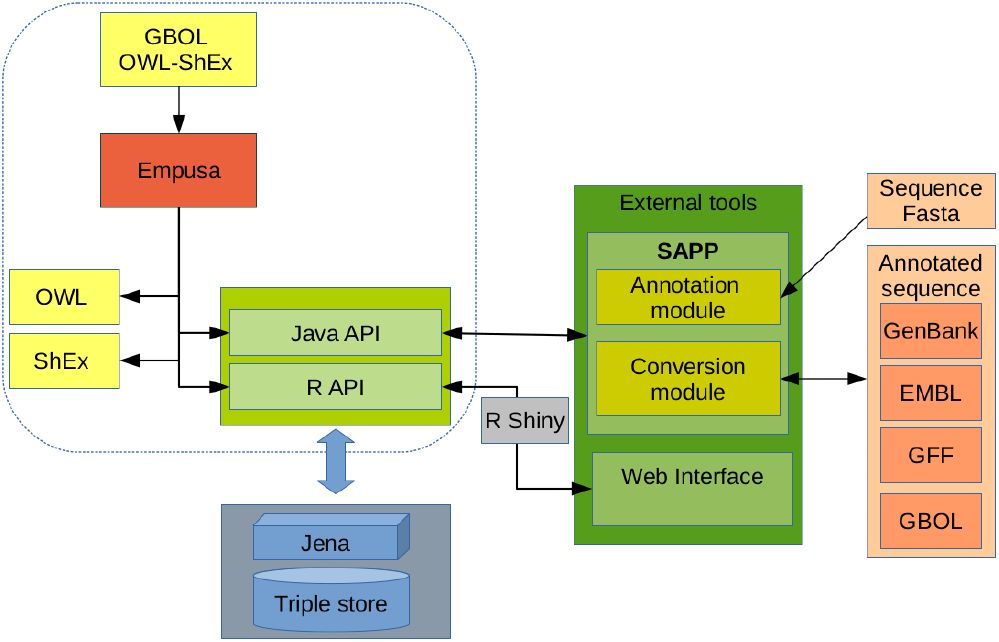
Schematic of an interoperable provenance centered genome annotation pipeline. The GBOL stack (dashed box) provides the Genome Biology Ontology Language (GBOL) (Yellow) and associated infrastructure to keep it consistent and extendable (Empusa). The SAPP module functions as an interface for (standardly used) genome annotation tools. Using the JAVA API, SAPP retrieves raw genome data from the triple store, runs genome annotation tools in batch and uses the GBOL ontology to automatically store their predictions and associated data provenance directly as RDF triples in the triple store database (Blue). Stored predicted functional annotations, data provenance and linked meta-data can be queried within JAVA and R with SPARQL and by using a web interface (Green). Parsers have been developed for conversion of annotation files in standardly used formats (Orange).

### Embedding with other ontologies

GBOL is embedded in the corpus of currently developed web technologies and when possible we have integrated existing ontologies such as: FALDO (13), PROV-O (22), SO (12), SBOL (14), BIBO (23), WikiData (24), FOAF (25), Gene ontology (GO) (26) and the Evidence ontology (27) as depicted in Figure 3. Annotation of genomic location is inspired by FALDO ontology, although several elements had to be modified. The PROV-O ontology was used and extended to store data provenance. Whenever applicable, we added a cross-link to exact matching terms within the FALDO, SO and SBOL ontologies. Identification of persons and institutions is done through the FOAF ontology and BIBO is used to identify publications. GBOL does not represent a vocabulary to describe genetic, molecular or cellular functions. Instead, terms can be cross-referenced to the many vocabularies that provide functional descriptions to the (products of) genetic elements, such as Gene Ontology, Enzyme commission (EC) numbers, and the CHEBI and RHEA databases (28, 29), among others.

**Fig. 3.**
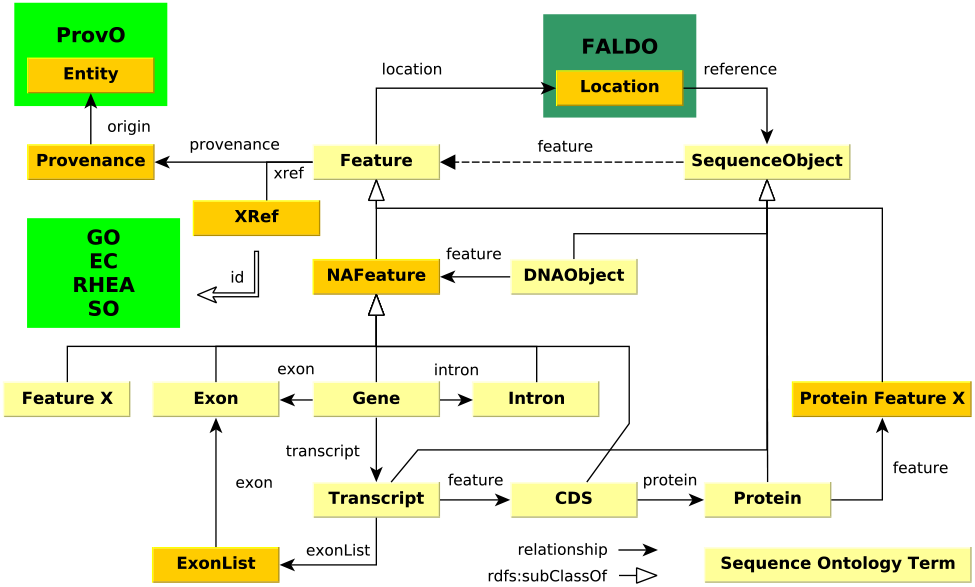
Embedding of the GBOL ontology with already existing ontologies. FALDO, ProvO, GO, EC, RHEA and SO are existing ontologies. Classes are in yellow and an explanation is provided in the main text.

### Key GBOL classes

Common elements in genome annotations include different classes of DNA molecules such as chromosomes, plasmids and contigs, genes, transcripts, exons, introns, proteins, protein domains and functional annotations. The following sections summarize the key classes of the ontology. An extensive description for each element can be found in the documentation available at http://gbol.life/0.1/.

#### Genomic locations

Genomic locations of all features in GBOL is captured with the *Location, Position* and *StrandPosition* classes, which are inspired by the FALDO ontology and represented in Figure 4. The *Location* and its subclasses together with the *StrandPosition* define an interval on the Sequence, whereas *Position* defines a single position in a sequence. A location can be either: i) A region which has begin and end positions; ii) A collection of regions (ordered or unordered); iii) A single base at a given position; or iv) an *In-Between* location denoting a location between two bases after the base of which the position is given. Each region, base and in-between location can be defined to be located on the forward, reverse or both strands, although no strand should be specified if the sequence is a single stranded DNA sequence or a protein sequence. It should be noted that elements of a collection of regions can be located on different sequences. This can be used to encode cases in which an otherwise indistinguishable genetic element is located on multiple chromosomes.

**Fig. 4.**
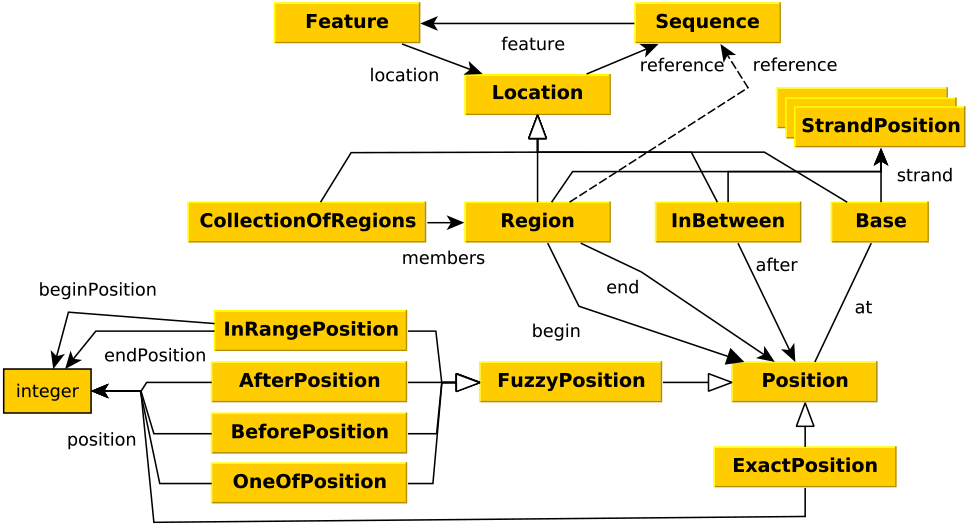
Graphical view of the GBOL ontology for genomic locations. An explanation of the classes is provided in the main text.

Exactly known positions can be indicated using the *Exact-Position* class containing the *position* property. Otherwise a not exactly known position, also called fuzzy position, can be indicated using either the *BeforePosition* class containing the *position* property, the *AfterPosition* class containing the *position* property, the *InRangePosition* class containing the *beginPosition* and *endPosition* properties or the *OneOfPosition* class containing multiple *position* properties.

#### Genes, transcripts and other commonly encountered genomic features

GBOL has a consistent model for storing genes, exons, (alternatively spliced) transcripts, coding sequences and proteins. Central to this model is the *Sequence* class that can have multiple annotations represented in the *Feature* class. An overview is provided in Figure 5.

In GBOL a sequence can be specified as a nucleic acid (NA) or a protein sequence. The sequence is attached to the *Sequence* class via the *sequence* property, provided in the DNA, RNA or protein encoding standard. NA-sequences can represent transcripts or other elements such as chromosomes, plasmids, scaffolds, contigs or reads. No distinction is made between DNA and RNA and the *strandType* denotes that it is either a double or single stranded DNA or RNA. As indicated in Figure 5 the type of sequence determines the features it might be associated to (*ProteinFeature, NAFeature* or *Tran-scriptFeature*),

**Fig. 5.**
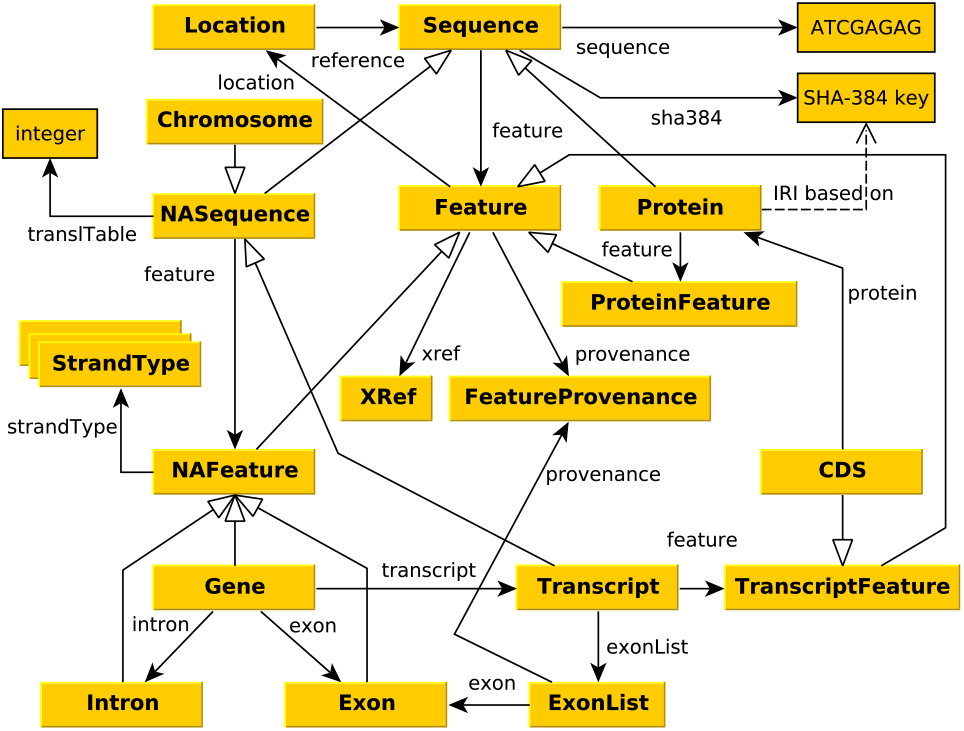
Graphical view of the GBOL ontology for genes, transcripts and other commonly encountered genomic features. An explanation of the classes is provided in the main text

Typically, each *GBOLDocument* contains one or more *NASequences* (e.g. *Chromosome, Contig, mRNA),* which can have multiple features including all gene, exon, intron, sequence variations, and structural, regulatory and repeat annotations. Each gene is linked to its associated exons, introns and transcripts. Due to alternative splicing a gene can have multiple transcripts. Each transcript has its own unique list of exons, which is linked through the *exonList* and associated *exonList* class to all associated exons. A transcript can be either a mRNA, ncRNA, rRNA, tmRNA, tRNA, precursor RNA or a miscellaneous RNA. The type of transcript determines the associated features: mRNA transcripts can have features linked to coding sequence (CDS), 5’-UTR, 3’-UTR and poly A tail. The mRNA translation table is defined with the *translTable* property from the parent sequence. The association between CDS and the encoded protein is preserved and information about the translation is stored if it is different from the default translation (for example, use of alternative stop codons). Each protein has a unique IRI (http://gbol.life/0.1/protein/<SHA-384>) based on the SHA-384 hash of its sequence. This makes it possible to combine protein information from heterogeneous sources, as a protein can be associated to several CDS features. All information related to the protein which is unique to the genome (such as location) should be stored in the CDS feature. Protein annotation features may include, among other, conserved regions, protein domains, binding sites, 3D structure, signal peptides, transmembrane regions, and immunoglobulin regions. Operons can be defined with the *Operon* feature, to which other genomic features, such as genes, can be associated. Additionally, viral genome integration can be denoted using the *IntegratedVirus* feature.

#### Provenance related classes

Three types of provenance can be distinguished. Metadata refers to the owners of the samples, the biological origin, culture conditions etc. Dataset- and element-wise provenance pertain to the annotation process. All data within a single data collection stored in GBOL is based on the *GBOLDataSet,* which holds among other, references to all included samples, sequences, organisms, annotation results and linked databases. An overview of the document structure is given in Figure 6.

**Fig. 6.**
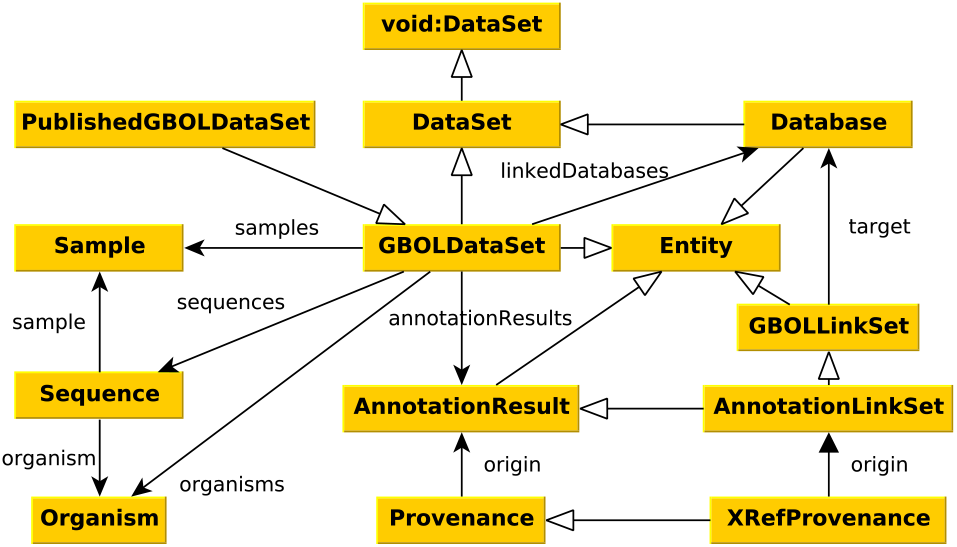
Graphical view of the GBOL Document structure. An explanation of the classes is provided in the main text

A sequence originates from a sample and samples are related to one or multiple organisms. The *sample* property which links to the *Sample* class describes where, when, how, by whom and from what the sample was collected. The fields follow the GenBank format. The *organism* property describes the taxonomic reference, its scientific name and its taxonomic lineage.

All annotations made within the *GBOLDataSet* have associated provenance and should originate from one of the listed annotation results, so that correspondence with originating databases is preserved. The *Database* and the *GBOLDataSet* classes are both sub classed from the void ontology, *Dataset* class contains a general description, including among other title, description, comment, license, version, data download address, SPARQL endpoint URI, and URL encoding.

#### Dataset-wise provenance

Storage of the dataset-wise provenance is based on the PROV-O ontology in which the *Entity, Agent* and *Activity* classes are central. An activity can use and generate entities, which are executed (*wasAssociated-With*) by an agent. As a result, an entity can be attributed to an agent. The GBOLDataset, AnnotationResult, *GBOLLinkSet* and *Database* classes (indicated in Figure 6 and 7) are subclasses from the PROV-O ontology *Entity* class, so that for each of these objects provenance on how, when and by whom they were created can be associated.

**Fig. 7.**
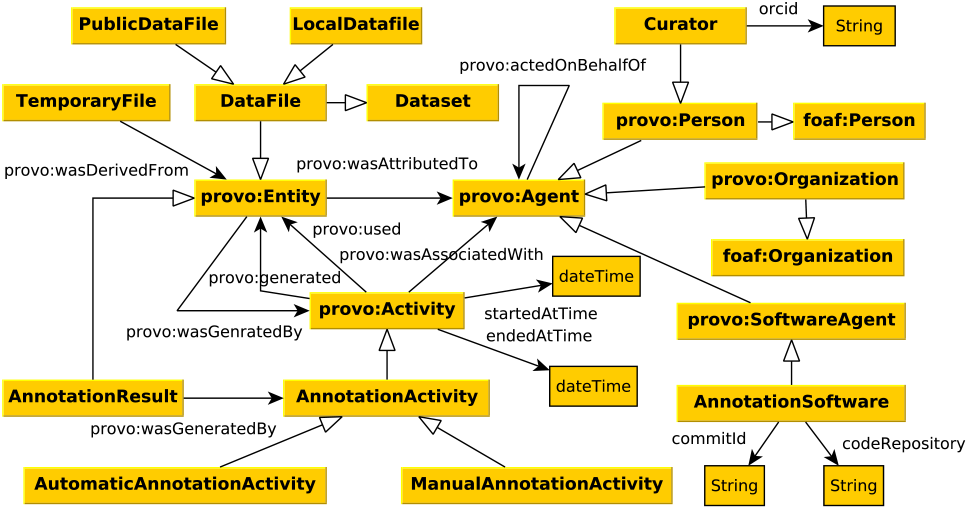
Graphical view of the GBOL Dataset-wise provenance. An explanation of the classes is provided in the main text

In GBOL an *Entity* is either a file or an annotation result. The annotation result is a set of triples contained within a GBOL document, whereas a file represents a physical file either on a computer or network. An *agent* can either be a curator, person, organization or annotation software. For the annotation software a version and code repository with associated commit identifier is included to enable univocal identification. For a curator, an ORCID (30) must be specified so that each curator can be uniquely identified together with his/her organization. Both *Person* and *Organization* are sub-classed from the FOAF ontology to include additional information such as name and email address.

Within GBOL, each activity is an annotation activity, which can be either an automatic process or a manual curation activity, with a start and end time. An automatic annotation must be associated with a software agent and the set of parameters used must be specified including the corresponding input and/or output files. Finally, manual curation must be associated with a curator.

#### Element-wise provenance and qualifiers

In addition to the dataset-wise provenance, GBOL is able to capture an additional layer of element-wise provenance, as the provenance of all the annotation in GBOL is captured per property per feature with the *FeatureProvenance,* as shown in Figure 8. For properties that could have items from multiple sources, we have defined the *Qualifiers,* each with its own associated provenance. A qualifier can either be a *citation, note* or cross reference (indicated by *xref).* A citation can hold a reference to literature encoded with the BIBO ontology.

**Fig. 8.**
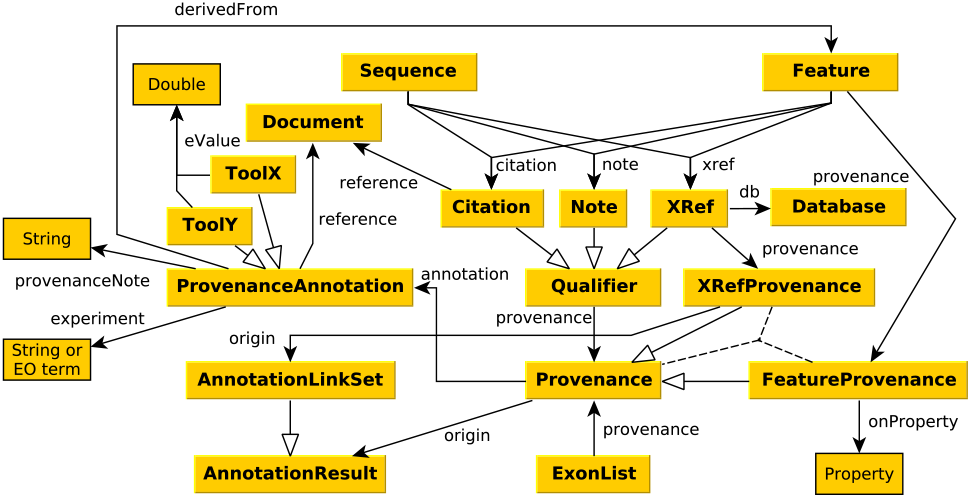
Graphical view of the GBOL element-wise provenance. An explanation of the classes is provided in the main text

Annotations are linked to the provenance object either through the *provenance* property of the qualifiers or the *onProperty* property of the *Provenance* feature. The provenance object links to both the dataset-wise provenance and the element-wise provenance. The *origin* links the provenance with the dataset-wise provenance (*AnnotationResult*), which includes among other the creation time, identity of the creating agent and the used parameters, as previously mentioned. The *annotation* links to the element-wise provenance (*ProvenanceAnnotation*), which includes: A free text note to describe the annotation; A list of references supporting the note; An experimental code, preferably from the Evidence Ontology to qualify the evidence supporting the conclusion; An optional *derivedFrom* that links to other features on which it is based.

Finally, each annotation tool generates its own evidence statements, often embedded in a statistical framework, characteristic of the algorithmic approach taken, such as p-values, bit scores, matching regions or any other scoring system. To store tool specific confidence scores, a subclass of the *ProvenanceAnnotation* class can be created. Some example classes include *Blast*, *HMM* and *SignalP* associated with the output of corresponding tools (31–33) However, these classes are not part of the GBOL ontology itself.

### Empusa

During the development of the standard, difficulties were encountered in managing the large set of properties and structures in the OWL and ShEx definitions and the API needed to encode the annotation information in conjunction with the associated provenance. Moreover, Analyses of various public repositories have shown that inconsistent, non-enforced usage of ontologies leads to mismatches between the descriptive OWL file and the actual content (15). In order to shorten the development cycle and to maintain consistency within and between the OWL and ShEx definitions and the API, a standalone tool was developed named Empusa. The input definition of Empusa is a combination between OWL and a simplified version of ShEx, which can be edited within Protégé (34). The classes are defined in OWL, whereas the properties are defined in each class under the annotation property ‘propertyDefinitions’ encoded within a simplified format of the ShEx standard. Additionally predefined value sets (for example all regulatory types) can be defined by adding a subclass to the EnumeratedValueClass. Each subclass of the value set is represented as one element within the value set. As standalone tool, Empusa can automatically and consistently generate an OWL and a ShEx definition, ontology documentation in markdown, an API, a JSON-LD framing file and a visualization. Empusa uses parts of the RDF2Graph tool (15) to generate a representation that can be subsequently used to generate a visualization within Cytoscape (35). This allows users to browse the complete ontology intuitively.

## Discussion

Comparative genome analysis is essential to understand the mechanisms underlying evolution and adaptation. Ideally, comparative genomics should be performed at the functional level, as this is highly scalable and more resistant to phylogenetic distances (36). However, as functional annotation is performed in a non consistent manner the current practical level of interoperability is at the sequence level. Many tools exists to obtain orthologous clusters which are shaped by a generalised acceptance threshold for similarity and alignment length which is a trade-off between sensitivity and false discovery (37, 38). At large scales these analysis are hampered by the high computational cost for finding bi-directional best matches. We have shown (36) that functional comparison, based on consistently annotated protein domains, provides a fast, efficient and scalable alternative.

The prerequisite of a direct comparative functional analysis is consistent annotation of the genetic elements with evidence statements. Recording the provenance allows class-specific cut-offs for each individual annotation. Element-wise provenance enhances the re-usability of the annotations, and allows the development of methods to combine evidence statements, often derived from complex statistical frameworks, into confidence statements. Element-wise provenance also enables a quick re-evaluation of evidence, for instance by using a tunable cut-off score.

GBOL has been developed to explore available genome sequences using the mining possibilities of linked data. As a result, GBOL has evolved to consistently capture annotation data generated by the Semantic Annotation Platform with Provenance (SAPP), available at http://semantics.systemsbiology.nl. Previous versions of the GBOL ontology have been used to compare 432 *Pseudomonas* strains through integration of genomic, functional, metabolic and expression data (39). Here GBOL was essential to capture, store and interlink the genomic and functional annotation data. Strikingly, over 432 *Pseudomonas* strains, consistent *de-novo* annotation yielded 838 additional GO-terms and 146 additional protein domains which would not have been identified using the original gene predictions. In addition to determining the functional pan- and core genome of a species, comparative genomics also enables the investigation of genotype-phenotype associations. In (40) we consistently functionally annotated and compared 80 publicly available mycoplasma genomes. The resulting semantic framework allowed us to efficiently query for functional differentiation of various mycoplasma species in relation to host specificity and phylogenetic distance.

Consistent functional annotation within a semantic framework requires a standardised ontology for the annotated elements and the associated based-on provenance. Linked data ensures that queries can be performed, mining multiple sequences at once, thereby providing a scalable alternative for large scale genome comparisons. The GBOL stack provides the ontology and corresponding API that enables the incorporation of functional annotation and provenance reducing complexity and is the outcome of efforts in a number of studies related to functional comparative genomics. Currently the GBOL stack is being used in various collaborative projects to handle genomic data of organisms across all domains of life (41–44).

GBOL has been primarily designed to handle genomic annotation. However, it has been designed in a modular and extensible manner so that in the future it can be extended to host other omics data types as proteomics and transcriptomics. The modular design of GBOL ensures that other ontologies can be incorporated and managed as separate entities. For instance, the majority of the feature and sequence classes within GBOL can be connected with those from the Sequence Ontology and are therefore linked with the *skos:exactMatch* predicate. The major difference between GBOL and SO is that SO has been defined as vocabulary of terms related to genetic elements, whereas the GBOL classes have been designed to describe genetic annotation and elements located on a sequence and is inspired on the principles of the GenBank format. However, still a number of features in the SO are not currently available in GBOL and future work should focus on including them. Another possible extension would be to link to other Minimum Information Standards like MIGS and extensions thereof (MIMARKS, MIxS) (45,46) and cross domain experiment reporting standards like ISA-tab (47). Other possible extensions relate to the development of the subontologies GBOL links to. For instance, BIBO is used to store information on literature references, however the OWL ontology file of BIBO has to be further improved, as it does not specify to which classes all of the properties should belong. Therefore we have chosen to include a less consistent representation of the properties by adding all properties to the root class *bibo:Document*.

Empusa, a core part of the GBOL stack, ensures the correct usage of the ontology through the provided R and JAVA API. We have ensured that Empusa can be used independently of GBOL (documentation available at http://gbol.life) and therefore can be used to develop new ontologies combined with an automatically generated API and documentation. This reduces the complexity and time to extend and develop ontologies with corresponding API’s and ensures consistent and correct usage of a defined ontology.

## Conclusions

Large scale analysis of heterogeneous biological data is hampered by lack of interoperability. To improve the exchange of information formalized information models are required. GBOL provides a formal representation of genomic entities, their properties and relations. The GBOL Stack provides a framework to enforce consistent and correct usage of GBOL. The semantic basis and the integration of provenance enables FAIR genome annotations, thereby unlocking the potential of functional genome annotation data.

## Methods

The GBOL ontology is OWL encoded and a ShEx schema is provided. All supporting software (Java and R API, Empusa) are written in Java with Gradle as build system. We use JENA (48) for handling and loading the RDF data into a triple store. Protégé was used for editing the ontology(34).

Storage of the genomic location is inspired by FALDO, although several elements had to be modified e.g. to account for features that start and end on different sequences. Differences include: i) *StrandPosition* is not subclassed from *Position*. Instead, an additional property is added to the region, *base* and *InBetween* location, this is done because these location object types can have both a strand position and an index position on the sequence. ii) The *reference* property is not part of a Position, but of a Location, because a location that starts on one sequence and ends on another sequence is an undefined sequence. iii) The *BaseLocation* and the *InBetween-Location* classes have been added to the ontology. iv) The *BaseLocation*, *InBetweenLocation*, *CollectionOfRegions* and *Region* are children of the *Location* class, such that the rest of the ontology can incorporate these classes. v) The *before* and *after* positions have been explicitly defined to include their semantics. vi) The classes sub-classed from *FuzzyPosition* have an integer to denote the position and do not point to another position object, which could allow for arbitrary complex location denotations. vii) The N- and C-terminal positions have been removed and all indexes are counted from the N-terminal side. Counting from the C-terminal side can be calculated based on the sequence length. vii) The reflective properties *beginOf* and *endOf* have been removed, because a position can also be referenced by the added base location. For consistency we have redefined all FALDO elements within our own namespace.

Cross-links to exact matching terms from other ontologies (such as FALDO, SO and SBOL) where added using skos:exactMatch. Additionally, several properties within the ontology point to existing ontologies, for instance: i) The *signalTarget* property of SignalPeptide, the *modification-Function* of *ModifiedResidue* and the *organelle* of *Sample* are interlinked with GO terms. ii) The *experiment* property of ProvenanceAnnotation, which denotes upon which evidence the annotation is based on, should point, where possible, to a term within the Evidence Ontology. iii) The *residue* property of *ModifiedResidue* must point to a term within the Protein Modification Ontology (49). iv) GBOL includes the GO terms for *tissueType* of the Sample class and points, when possible, to a term within the BRENDA Tissue and Enzyme Source Ontology (50).

The source file of the ontology encoded in the Empusa and associated generated OWL definition, ShEx schema and visualization for Cytoscape available at http://www.gitlab.com/GBOL under the MIT license. The generated Java and R API are available at https://gitlab.com/gbol/GBOLapi and https://gitlab.com/gbol/RGBOLApi under the MIT license. The conversion module, which is part of SAPP, is available at http://www.gitlab.com/SAPP/conversion under the MIT license. The supporting Empusa code generator is available at http://www.gitlab.com/Empusa under the MIT license. All projects are coded in Java and are based on the Gradle build system. All terms are resolvable and can be browsed for at the associated website http://gbol.life/0.1/.

## ACKNOWLEDGEMENTS

We thank Benoit Carreres for helpful design discussions. This work has received funding from the Research Council of Norway, No. 248792 (DigiSal) and from the European Union FP7 and H2020 under grant agreements No. 305340 (INFECT), No. 635536 (EmPowerPutida) and No. 634940 (MycoSynVac).

